# PACMOS: an R package for Projection And Classification of Multi-Omic Samples

**DOI:** 10.64898/2026.07.01.735542

**Authors:** Lipika Kalson, Alexandra Sexton-Oates, Gabrielle Drevet, Lynnette Fernandez-Cuesta, Matthieu Foll, Nicolas Alcala

## Abstract

**Motivation:** Integrated multi-omic analyses have transformed our understanding of cancer biology, giving rise to data-driven molecular classifications that capture disease heterogeneity beyond conventional histopathology. Among these approaches, multi-omic factor analysis (MOFA), a multimodal extension of principal component analysis, has been widely used to identify sources of molecular variation across omic layers and classify samples into molecular groups. However, classifying query samples according to an existing MOFA-based classification remains challenging, as there is no validated computational method for projecting samples into pretrained MOFA latent factor spaces.

**Results:** We present PACMOS, an R package that provides a generalizable approach to project query samples into pretrained MOFA latent factor spaces. We validate PACMOS using two cancer datasets with published MOFA-based classifications—lung neuroendocrine neoplasms and pleural mesothelioma—showing that PACMOS preserves the existing MOFA latent factor space while allowing query samples to be classified.

**Availability and implementation:** PACMOS is an open-source R package available at https://github.com/IARCbioinfo/PACMOS and archived on Zenodo at https://doi.org/10.5281/zenodo.20933824, along with installation instructions and a vignette.

**Supplementary information:** Supplementary data are available in separate files.

**Key messages:**

- PACMOS enables the projection of query cancer samples into pretrained multi-omic latent factor spaces.
- The package supports both continuous and discrete classifications.
- PACMOS provides a reproducible, per-sample workflow implemented in an R package.
- PACMOS demonstrates robust performance on pleural mesothelioma and lung neuroendocrine tumor datasets.

## 1 Introduction

Cancer classification remains a central challenge in oncology, driven by extensive molecular heterogeneity both across and within patients (Pe’er et al., 2021; Dagogo-Jack and Shaw, 2018). Traditional classifications are often based on histopathology or single-omic profiling—explaining only a small fraction of inter-patient variance, limiting the ability to explain clinical variability and guide precision treatment strategies (Drevet et al., 2026; Abdelaziz et al., 2024; Karczewski and Snyder, 2018).

Multi-omic analyses have transformed our capacity to characterize this heterogeneity by integrating multiple molecular layers (e.g., genomic, transcriptomic, epigenomic, proteomic) to obtain a comprehensive view of tumor biology (Hasin et al., 2017). Among these approaches, multiomic factor analysis (MOFA; Argelaguet et al. 2018, 2020)—a multimodal extension of principal component analysis (PCA)—has become a widely adopted standard for unsupervised classification. While current WHO classifications remain primarily grounded in histopathological criteria, multi-omic frameworks can provide complementary biological insights that can inform future classification efforts (WHO Classification of Tumours Editorial Board, 2021). For example, the MESOMICS study showed that MOFA captured ∼ 60% of inter-patient molecular variance in pleural mesothelioma, compared with *<* 10% explained by the current WHO classification based on cell morphology, thereby providing a far more informative description of the disease (Mangiante et al., 2023). Similar applications of MOFA in lung neuroendocrine tumors have supported the identification of molecular groups with potential relevance for prognosis and treatment (Alcala et al., 2019; Gabriel et al., 2020; Leunissen et al., 2025; Sexton-Oates et al., In press), and multi-omic profiling of clear cell renal cell carcinoma (Hu et al., 2024) and breast cancer (Sharma et al., 2024) have also revealed subtypes of clinical interest. Multi-omic approaches have also been applied to deepen the understanding of biological processes underlying carcinogenesis without necessarily proposing new classifications, as illustrated in kidney cancer where MOFA linked aging, epithelial-mesenchymal transition, and xenobiotic metabolism to tumor progression (Penha et al., 2024).

Despite their value, current multi-omic molecular classifications remain primarily discovery-oriented, with limited reuse in independent datasets due to challenges in standardization, reproducibility, and prospective validation (Mohr et al., 2024; Akhoundova and Rubin, 2022). In particular, published frameworks often define a MOFA latent factor space or molecular groups within a reference cohort, but provide limited practical guidance for projecting query samples into the same space (Mangiante et al., 2023; Sharma et al., 2024; Hu et al., 2024; Sexton-Oates et al., In press). This is mostly due to the fact that MOFA is not a simple matrix factorization but a probabilistic Bayesian model with omics-specific weight matrices (Argelaguet et al., 2018), so query sample projections cannot be obtained by simple matrix multiplications as for PCA. Only a few recent efforts have addressed this issue, such as models developed to classify non-TCGA cancer samples into established TCGA molecular subtypes using compact feature sets (Ellrott et al., 2025), but they do not provide a general method for MOFA-based classifications.

To address this gap, we present PACMOS, an R package that provides a reproducible and generalizable workflow for projecting query samples into pretrained MOFA-based latent factor spaces and performs continuous and discrete molecular classifications. PACMOS preserves the structure of the pretrained space and provides quality-control of the projection accuracy. While we illustrate its use on pleural mesothelioma and lung neuroendocrine tumors—two entities in which multi-omic studies have proposed clinically relevant classifications that have been discussed in the context of WHO classification refinement— PACMOS is designed to be broadly applicable to any MOFA-based classification.

## 2 Methods

PACMOS projects query samples into a pretrained MOFA latent factor space by sequentially adding each query sample to the pretrained MOFA, retraining the model, and checking that retraining did not alter the latent factor space. The projected samples are then assigned to molecular groups.

### 2.1 Data preparation

PACMOS requires three inputs. (i) *RData* objects providing the input matrices used for training MOFA in the reference cohort typically includes DNA methylation and additional omic layers such as genomic alterations or gene expression. (ii) Reference latent factor matrix. (iii) Multi-omic data from query samples. The query samples must contain at least one of the omic layer used in the reference cohort, and must have been processed exactly the same way: from pre-processing (e.g., mapping, variant calling) to normalization and scaling for continuous variables (e.g., quantile or variance-stabilization transformation for transcriptomics).

### 2.2 Sample-wise sequential integration

Each query sample is individually appended to the reference MOFA input matrices as a new column, with rows matching the order of features in the reference matrix; unavailable layers are encoded as a new column full of missing values. This generates one augmented dataset per query sample for independent model retraining.

### 2.3 Projection into the MOFA latent factor space

For each augmented dataset, a MOFA model is retrained using the same hyper-parameters and model structure as the reference model. The retraining step produces updated factor values for all samples, including the newly added query sample. To ensure comparability, latent factors are matched and sign-corrected to the reference latent factors using correlation based assignment with the Hungarian algorithm (Kuhn, 1955). An augmented latent factor matrix is then constructed by retaining reference latent factor values for all reference samples and inserting only the matched query sample factor values. Detailed latent factor matching and quality-control procedures are provided in the Supplementary document.

### 2.4 Classification

Following projection into the MOFA latent factor space, samples are classified using clustering approaches applied to the augmented latent factor matrix, including hard clustering—where each sample is given a unique label—and fuzzy clustering—where samples have multiple labels, each with a certain proportion of association with each group.

#### Hard clustering into molecular groups

For hard clustering, the k-means algorithm is applied, using the number of clusters and augmented latent factor matrix as inputs and multiple initializations to ensure robustness. In order to assign to each cluster a molecular group label (e.g. corresponding to a known molecular subtype), we assess the correspondence between reference samples and k-means clusters, assigning to each k-means cluster the reference label showing the maximum overlap with its members.

#### Fuzzy clustering with archetypal analysis

For fuzzy clustering, samples are modeled as convex combinations of predefined archetypes representing extreme molecular phenotypes (Cutler and Breiman, 1994). Archetype coordinates are defined *a priori* from the reference classification in the selected MOFA latent factor space. For each sample, archetype proportions were inferred using constrained least squares problem that minimized the squared Euclidean distance between the sample coordinates and a weighted sum of archetype coordinates. The inferred archetype proportions were constrained to be non-negative and to sum to one, allowing each sample to be represented as a continuous mixture of reference archetypes. The optimization was implemented in R using *limSolve::lsei*.

### 2.5 Applications

PACMOS is evaluated in two cancer datasets for which MOFA-based molecular classifications have been established.

#### 2.5.1 Lung neuroendocrine tumors

The lungNENomics cohort (*n*=319), (Sexton-Oates et al., In press) was used as a reference cohort. It includes multi-omic profiles along with multi-regional sequencing of 41 patients and represents the most comprehensive multi-omic dataset for this tumor type. The original study defined a MOFA-based latent factor space (Factors 1, 2, and 5) and a discrete classification into four molecular groups (namely, Ca A1, Ca A2, Ca B, and supra-carcinoid enriched).

PACMOS was applied to an external lungNET dataset from (Miyanaga et al., 2020), preprocessed consistently with the lungNENomics reference dataset (Kalson et al., 2026). Samples were projected into the lungNENomics MOFA latent factor space; a detailed workflow to perform the analysis is provided as a vignette in the PACMOS R package using two representative query samples.

#### 2.5.2 Pleural mesothelioma

The MESOMICS cohort (*n*=120, Mangiante et al. 2023) was used as the reference cohort for pleural mesothelioma. This dataset comprises multi-omic profiles from chemonaive, surgically resected tumors with matched clinical annotations. The original study defined a MOFA-based latent factor space driven by two latent factors (Morphology and Adaptive-Immune response) and a classification into three archetypes (Acinar, Tumor-immune interaction and Cell Division) using fuzzy clustering.

To illustrate PACMOS on query samples from an independent cohort, we applied PACMOS to pleural mesothelioma samples from the National Cancer Institute (Nair et al., 2023), preprocessed consistently with MESOMICS (Di Genova et al., 2023). A step-by-step vignette in the PACMOS R package illustrates this workflow using two representative query samples.

### 2.6 Validation

To validate PACMOS, we employed a leave-one-out (LOO) approach on the MESOMICS and lungNENomics cohorts, whereby each sample was iteratively excluded from the reference cohort and treated as a query sample. For each held-out sample, the reference MOFA latent factor space was reconstructed using the remaining samples, the held-out sample was subsequently projected back into the space using PACMOS, and correlations between reference (“true”) and projected (“predicted”) factor values were computed to assess the accuracy of the projection. We also examined whether the downstream classification of projected samples was accurately recovered, using either discrete clustering for lungNENomics or fuzzy clustering for MESOMICS.

As the lungNENomics study also provides discrete molecular class labels, we used it to validate PACMOS in a hard-clustering framework. We first reproduced the original four-group molecular grouping by running k-means (default parameters, *k*= 4) on the same MOFA latent factors (Factor 1, 2, and 5) used in the lungNENomics study, and assigning each cluster to a known molecular group according to the maximal overlap of samples. We then used this clustering as the reference classification. The latent factor values of projected LOO samples showed high correlation with their original MOFA factor values (*r* = 0.997, 0.993 and 0.957 for Factor 1, 2, and 5 respectively; Supplementary Fig. S1). In order to assess the relative importance of each omic layer in allowing accurate projection of query samples, we repeated this validation analysis across different omics-layer combinations, removing omic layers from the query samples accordingly. Latent factor recovery remained high for most combinations that included expression and/or methylation data, whereas genomics alone showed poor recovery of the latent factor values, with projected factor values collapsing towards the center of the MOFA latent factor space (Supplementary Fig. S2). This behavior is consistent with the lower amount of information contained in the sparse binary genomic features relative to continuous omic features, especially in low mutational burden cancer types such as lung neuroendocrine tumors (Hoadley et al., 2018). We then assessed whether projected samples were assigned to the same molecular group as in the reference classification. Overall, PACMOS yielded a classification accuracy of 99.1% in this discrete clustering framework (Fig. 1e (top)). High molecular group classification accuracy was also observed across most omic-layer combinations, particularly those including expression and/or methylation data (accuracies of 98.9% for G.E, 99.0% for M.E, 98.2% for E alone, 98.4% for M alone and 97.4% for M.G) whereas genomics alone showed accuracy of 49% (G:Genomics, E: Expression, M:Methylation; Supplementary Fig. S3).

**Figure 1:**
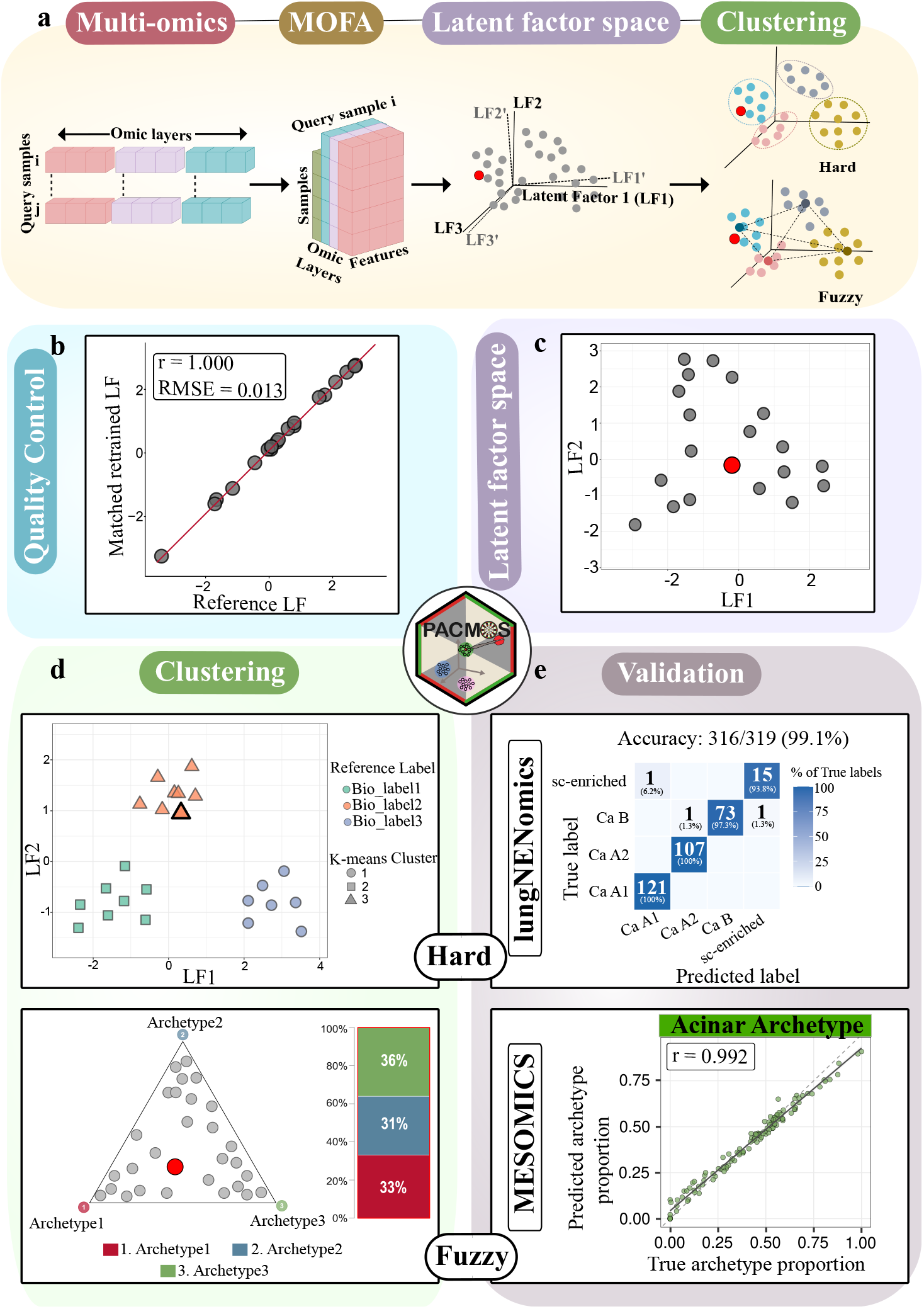
PACMOS overview and validation. (a) Overview of PACMOS. Multi-omic profiles from query samples are added one at a time to reference MOFA input matrices. The MOFA model is then retrained with these augmented inputs, outputting latent factor values for the query sample which can be assigned to molecular groups using either hard or fuzzy clustering. (b-d) Illustrative examples of PACMOS outputs; (b) MOFA quality control showing the correlation between reference factor values and their matched retrained factors values together with Pearson correlation coefficient (*r*), and RMSE values. (c) Query sample (red) projected into the reference latent factor space spanned by Latent factor 1 (LF1) and Latent factor 2 (LF2), highlighting its position relative to reference samples. (d) (top) Hard clustering: scatter plot of reference and query samples in the latent factor (LF1-LF2) space. The query sample is highlighted with a dark black border. Samples are colored by molecular group from the reference study, while shapes indicate k-means clusters. (bottom) Fuzzy clustering: all samples are positioned in a tetrahedron (triangle) whose vertices correspond to three archetypes and points are a convex combination of these archetypes, with the query sample in red; accompanied by a stacked barplot showing archetype proportions. (e) Validation using Leave One Out (LOO). (top) Hard clustering: confusion matrix showing classification predicted by PACMOS (x-axis) against true classification (y-axis). (bottom) Fuzzy clustering: scatter plot showing true archetype proportions (x-axis) vs PACMOS predicted archetype proportions (y-axis) across samples in the MESOMICS cohort.

For MESOMICS, we evaluated PACMOS in a fuzzy-clustering setting by assessing whether LOO projections matched each sample’s original factor values and its corresponding archetype proportions. The projected samples closely matched their original values along the two MOFA latent factors used in the MESOMICS framework, the Morphology and Adaptive-Immune response factors, with correlations of r=0.992 and 0.982 respectively (Supplementary Fig. S4). This concordance remained high across other omic-layer combinations, particularly when methylation and/or expression data were available, whereas projections based on genomics alone showed substantially weaker recovery of the factor values (Supplementary Fig. S5), as in the lungNENomics example. PACMOS also accurately recovered the archetype proportions of the three MESOMICS archetypes: Acinar (*r* = 0.992; Fig. 1e (bottom)), Cell Division and Tumor-immune interaction (r = 0.981 and 0.982 respectively; Supplementary Fig. S6). Across most-omic-layer combinations, archetype proportions remained highly correlated with the original reference values, particularly when transcriptomic and/or methylation data were available, while genomics alone again showed a low correlation across all three archetypes (Supplementary Fig. S7).

## 3 Implementation

PACMOS is implemented as an open-source R package (available via the IARC Bioinformatics GitHub, with Bioconductor submission in progress). The core functionality is organized into five main steps:

1. Construction of MOFA input matrices for each query sample (*s1_add_sample_to_mofa*); this requires precomputed MOFA input matrices and latent factors from a reference cohort, together with consistently preprocessed query multi-omic data. The workflow performs sample-wise integration by appending each query sample to the reference inputs.
2. Running MOFA on the input created above (*s2_run_mofa*);
3. Projection of query sample in reference latent factor space (*s3_plot_query_samples_mofa*);
4. Inferring archetype proportions or discrete clusters using either archetypal analysis or k-means clustering respectively (*infer_fuzzy_proportion* or *infer_kmeans_clusters*);
5. Visualization (*plot_fuzzy_query_sample* or *plot_kmeans_query_sample*).

Detailed documentation, tutorials and step-by-step examples are available in the package vignettes.

## 4 Conclusion

We provide an R package that allows the projection of query cancer samples into established multiomic molecular classifications. We show that PACMOS accommodates both continuous (fuzzy) and discrete (hard) classifications, validating its accuracy in two recently proposed molecular classifications: that of lung neuroendocrine tumors and that of pleural mesothelioma. Given that multi-omic molecular classifications are increasingly used to refine tumor classification and support clinically relevant patient stratification, PACMOS provides a timely and practical tool for researchers and clinicians seeking to use proposed molecular classifications and generate biologically and clinically relevant insights.

Note that PACMOS is currently specific to MOFA, as multi-omic integration methods do not use standardized inputs and do not provide standardized outputs. New PACMOS functions could be added in the future to handle other multi-omic integration methods, but each method will require a dedicated development and these extensions will thus need to be performed on a case-by-case basis.

## 7 Data availability

The data underlying this article are available in the R package PACMOS on IARC’s GitHub repository at https://github.com/IARCbioinfo/PACMOS and DOI in Zenodo: https://doi.org/10.5281/zenodo.20933824, and are based on previously published data from (Mangiante et al., 2023; Sexton-Oates et al., In press) also available in IARC’s github repository at https://github.com/IARCbioinfo/MESOMICS_data for the MESOMICS data and at https://github.com/IARCbioinfo/MS_lungNENomics for the lungNENomics data.

## 8 Author contributions statement

N.A. and M.F. conceptualized the study, L.K. performed the formal analyses, N.A., M.F., L.F.C, and L.K. acquired the funding, L.K. conducted the investigation and produced the software. L.K. and A.S-O. performed the validation and visualization, N.A., M.F., L.K. and G.D. created the methodology, N.A., M.F., and L.F.C supervised the work, N.A. and L.K. wrote the original draft. All authors reviewed and edited the manuscript.

## 9 Acknowledgments

The results published here are in part based upon data from the MESOMICS and lungNENomics projects, generated by the Rare Cancers Genomics initiative (www.rarecancersgenomics.com) led by the Computational Cancer Genomics team at the IARC (https://www.iarc.who.int/teams-ccg). It is also an output of the COALA network (https://coala-lung.org/).

