## Supplementary for "PACMOS: an R package for Projection And Classification of Multi-Omic Samples"

### Supplementary Figures

CONTENTS

Detailed Latent Factor matching method

Supplementary Figure S1

Supplementary Figure S2

Supplementary Figure S3

Supplementary Figure S4

Supplementary Figure S5

Supplementary Figure S6

Supplementary Figure S7

##### Detailed Latent Factor matching method

For each retrained MOFA model, latent factor matching is performed using the reference samples shared between original and retrained models. A Pearson correlation matrix is calculated between all latent factors inferred from retrained model and the selected latent factors from the reference models. To account for the arbitrary ordering and sign of MOFA latent factors during retraining, factor matching was formulated as a linear assignment problem. Specifically, a cost matrix was defined as (1- |*r*|^2), where r denotes the Pearson correlation coefficient between each pair of retrained and reference latent factors. The Hungarian algorithm was then applied to identify the one-to-one assignment that minimized the total cost, thereby selecting the combination of factors with the highest overall squared absolute correlation to the reference axes. Following assignment, factors with negative correlation to their matched reference latent factor were multiplied by (-1) to ensure consistent orientation.

Projection quality was assessed by calculating Pearson correlation and root mean square error between the original and matched latent factors across shared reference samples. PACMOS also generates factor-assignment summaries and diagnostic plots to verify recovery of latent factor structure.

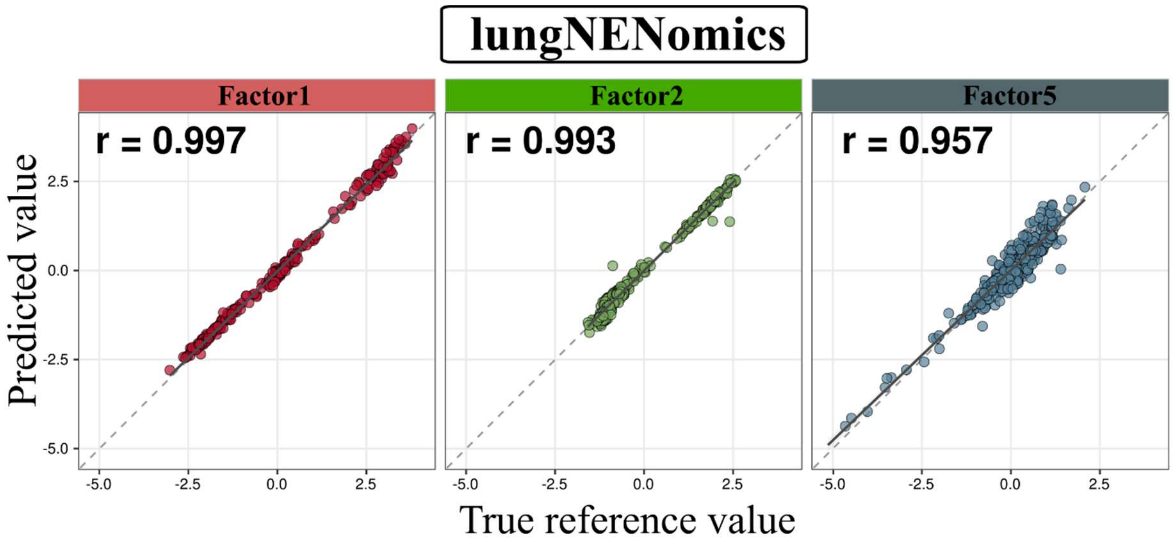


**Supplementary Figure S1. Leave-one-out validation of PACMOS: MOFA Latent Factors in the lungNENomics study.** Scatterplot comparing true MOFA latent factor values (x-axis) with PACMOS predicted MOFA latent factor values (y-axis) for each latent factor used in the lungNENomics study (Factor1, Factor2, Factor5). r: Pearson correlation coefficient.

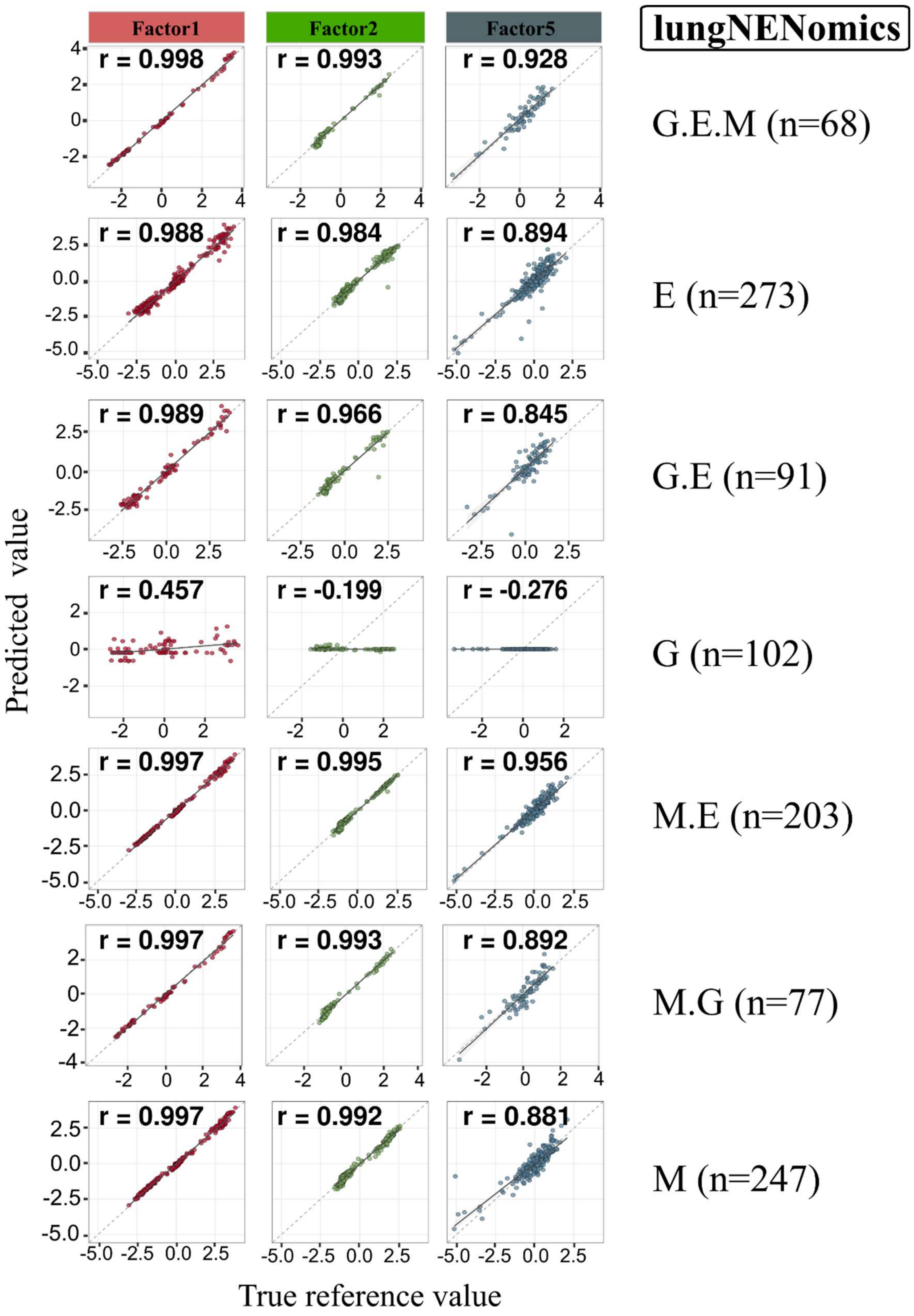


**Supplementary Figure S2. Leave-one-out validation of PACMOS: MOFA Latent Factors under different omic input combinations in the lungNENomics study.** Scatterplot comparing true MOFA latent factor values (x-axis) with PACMOS predicted MOFA latent factor values (y-axis) for each latent factor used in the lungNENomics study (Factor1, Factor2, Factor5). The right column indicates which omic layers were used as input (G: genome, E: expression, M: methylation). n: sample size. r: Pearson correlation coefficient.

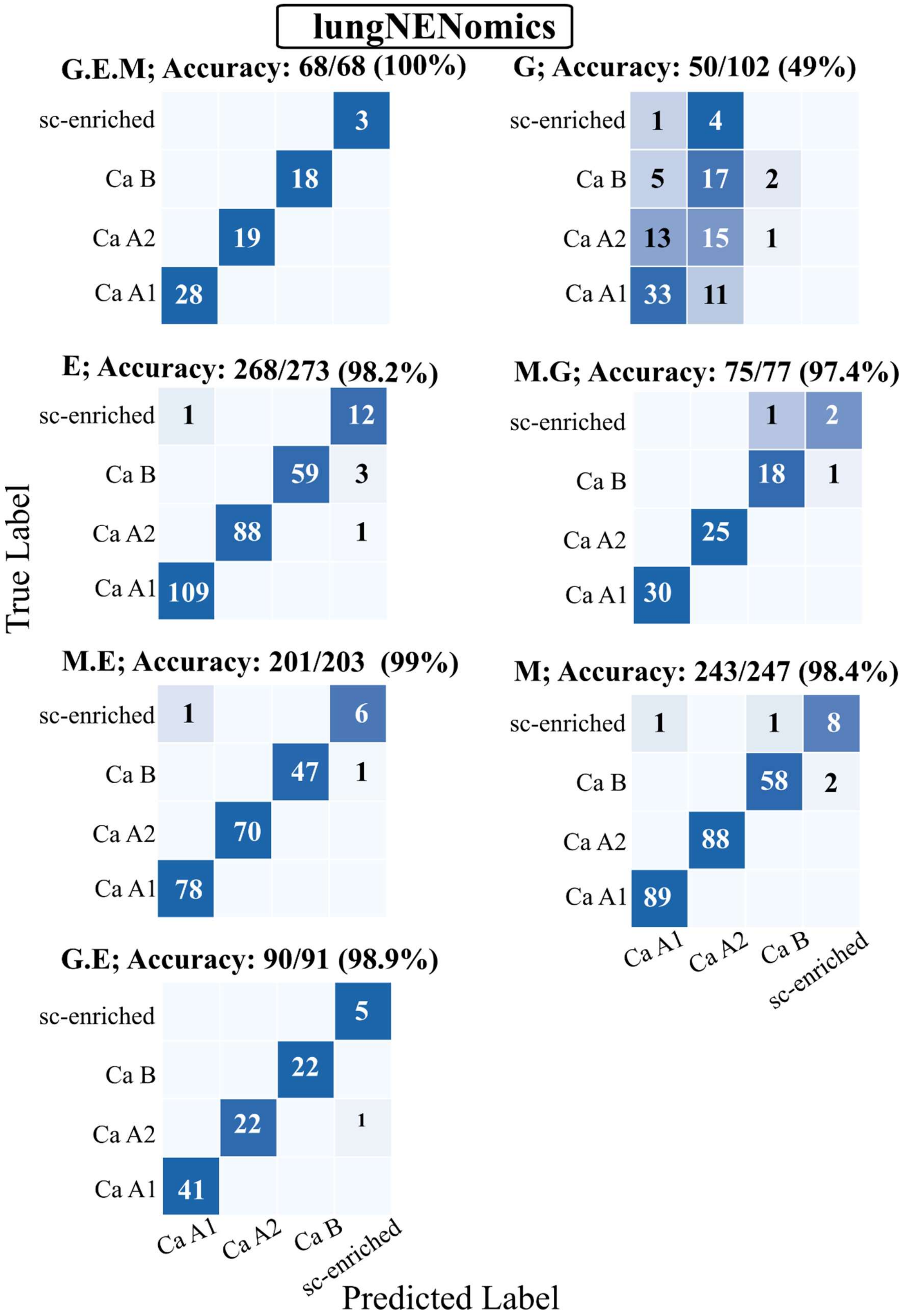


**Supplementary Figure S3**. **Leave-one-out validation of PACMOS: Hard clustering using different omic layers in the lungNENomics study.** Confusion matrices showing true (y-axis) and predicted (x-axis) molecular class using k-means clustering on projected MOFA factors (Factor1, Factor2, Factor5), for different combinations of omic laers (G: Genomic; M: Methylation, E: Expression).

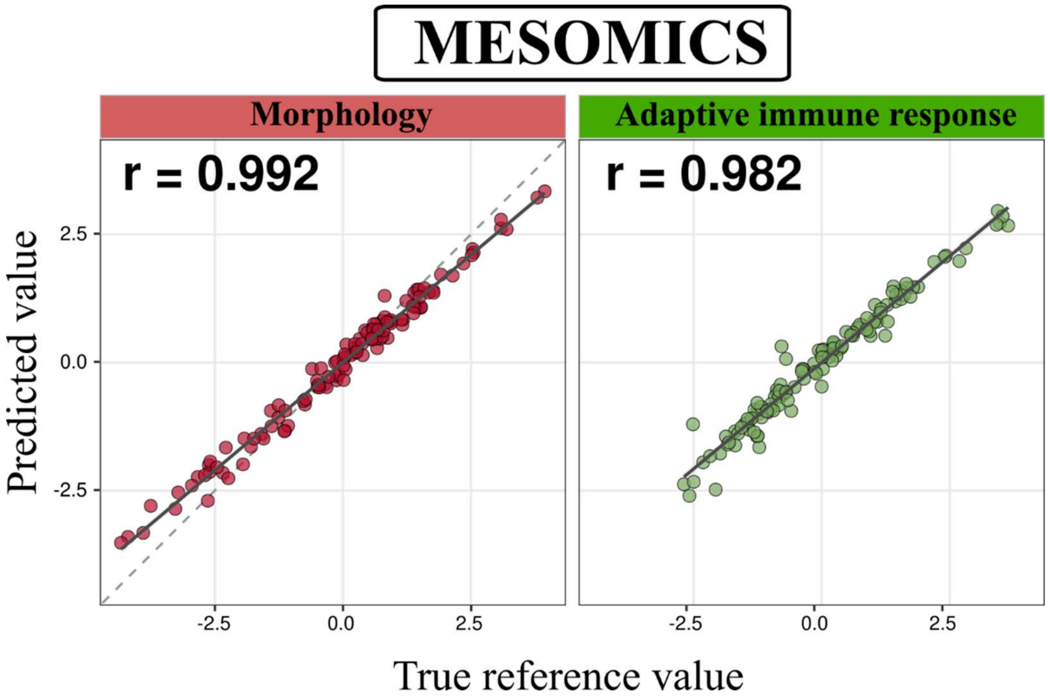


**Supplementary Figure S4. Leave-one-out validation of PACMOS: MOFA Latent Factors in the MESOMICS study.** Scatterplot comparing true MOFA latent factor values (x-axis) with PACMOS predicted MOFA latent factor values (y-axis) for each latent factor used in the MESOMICS study (Morphology and Adaptive immune response factors). r: Pearson correlation coefficient.

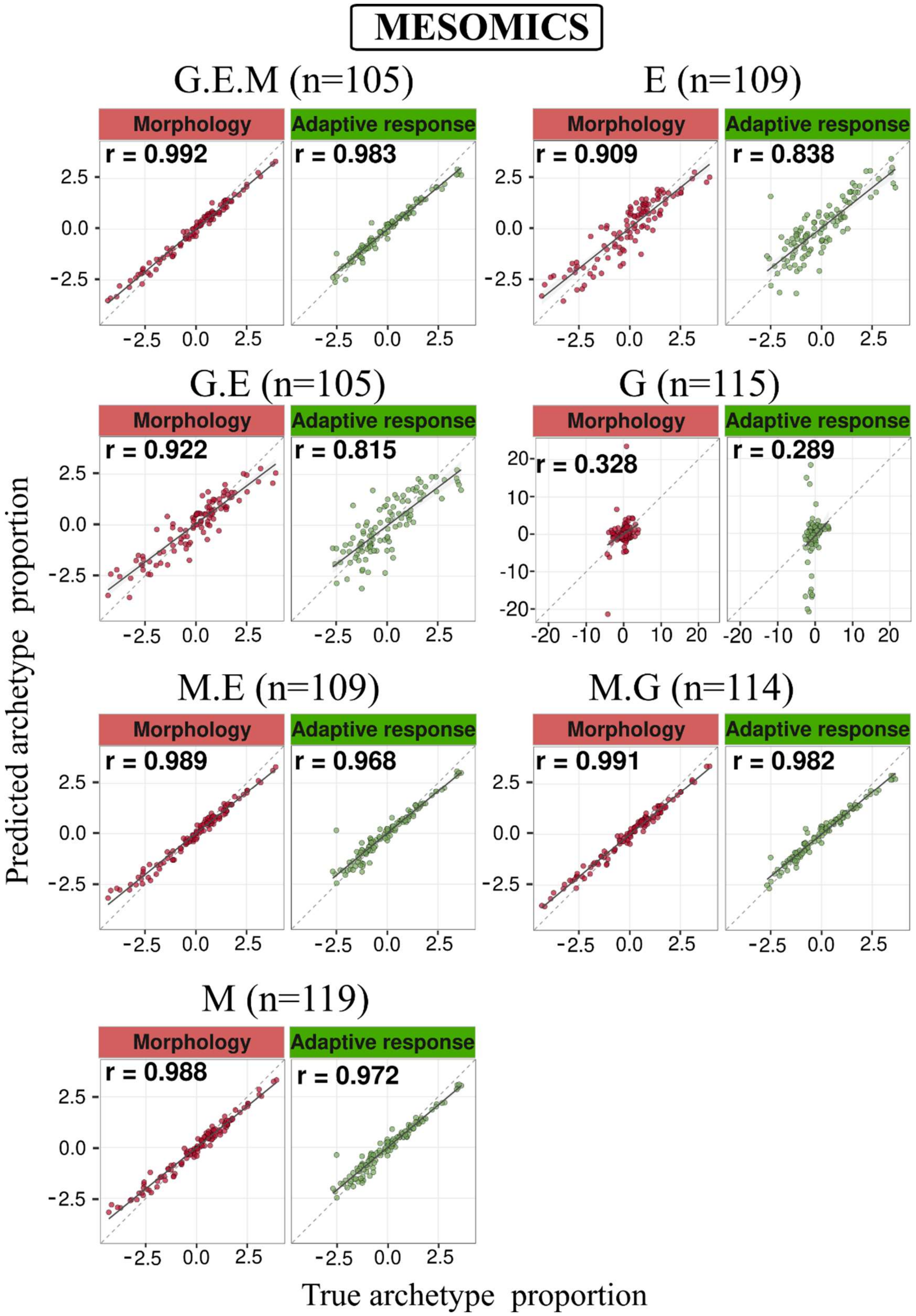


**Supplementary Figure S5. Leave-one-out validation of PACMOS: Fuzzy clustering using different omic layers in the MESOMICS study.** Scatter plot showing true (y-axis) and predicted (x-axis) molecular classes using k-means clustering on projected MOFA factors (Morphology and Adaptive immune response), for different omic layer combinations (G: Genomic; M: Methylation, E: Expression). r: Pearson correlation coefficient.

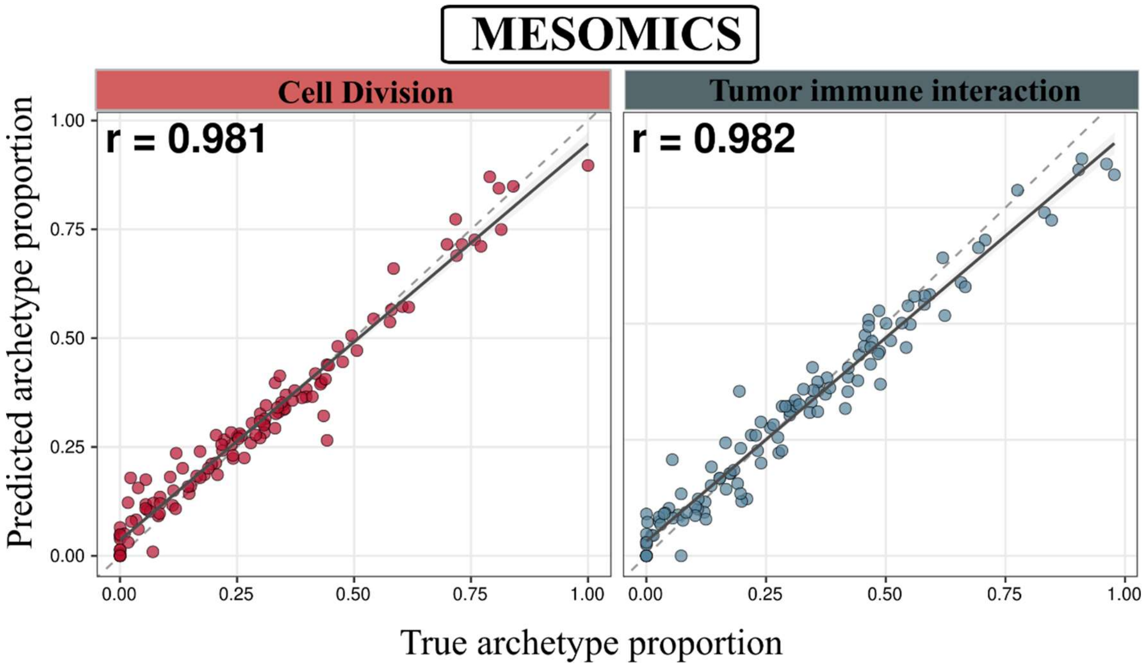


**Supplementary Figure S6. Leave-one-out validation of PACMOS: Archetype proportions in the MESOMICS study.** Scatterplot comparing true MOFA latent factor values (x-axis) with PACMOS predicted MOFA latent factor values (y-axis) for each latent factor used in the MESOMICS study (Morphology and Adaptive immune response factors). r: Pearson correlation coefficient.

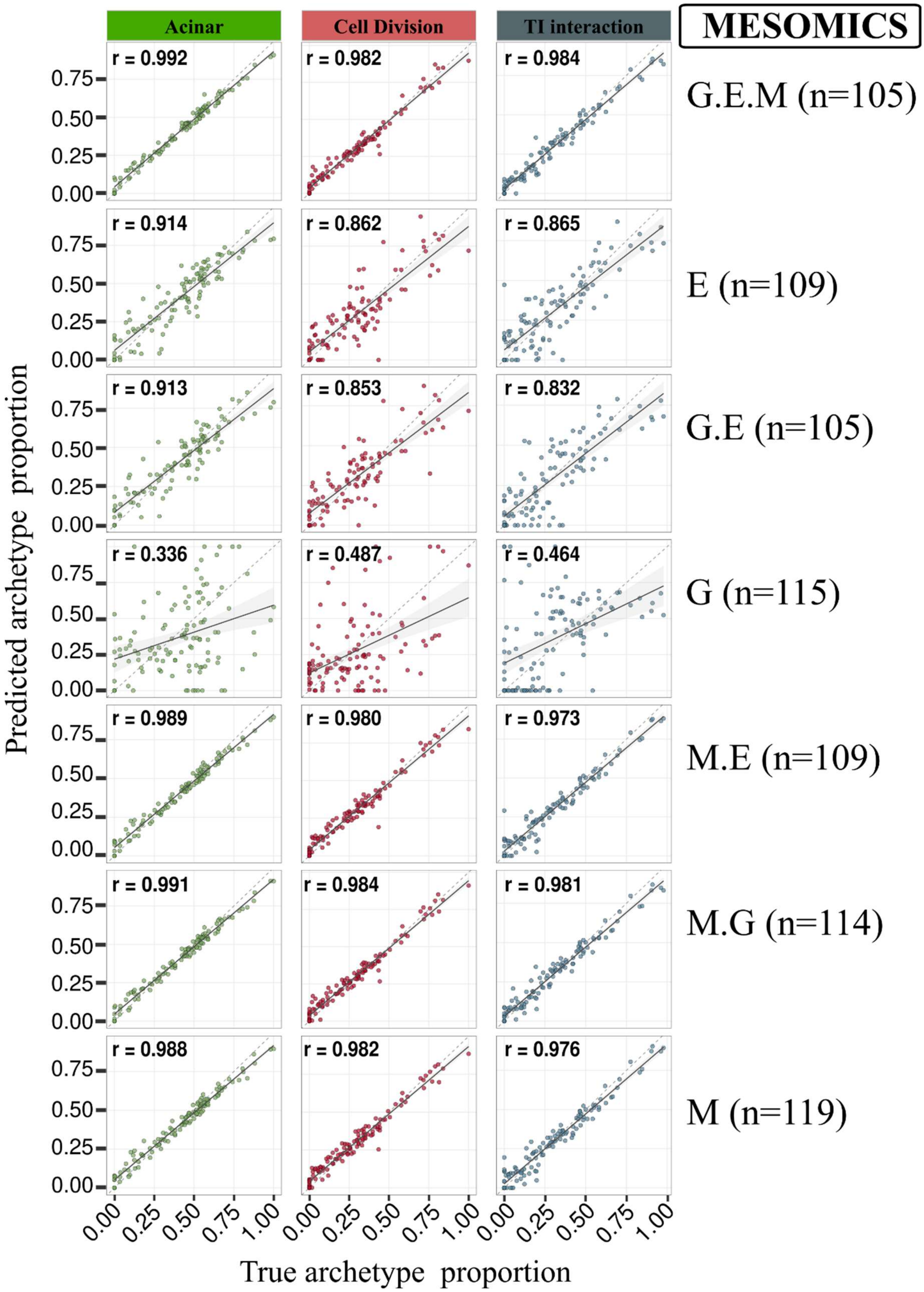


Supplementary Figure S7. **Leave-one-out validation of PACMOS: Fuzzy clustering using different omic layers in the MESOMICS study.** Scatterplot showing true (y-axis) and predicted (x-axis) archetype proportions on projected MOFA factors (Morphology and Adaptive immune response), for different combinations of omic layers (G: Genomic; M: Methylation, E: Expression). n: sample size. r: Pearson correlation coefficient.
